# Neutrophil and NET-driven pulmonary microvascular injury following myocardial injury: attenuation by S100A8/A9 inhibition

**DOI:** 10.1101/2025.09.12.675647

**Authors:** Danielle Lezama, Orestis Katsoulis, Eloise Marriott, Beata Grygielska, Dean Kavanagh, Celine Hsi Chen, Millie M Jackson, Ellen Jenkins, Katie Spencer, Emmanuel Abimbola, Nawaal Kiwia, Rahul Mahida, Davor Pavlovic, Helen M. McGettrick, Babu Naidu, Alexandru Schiopu, David R Thickett, Aaron Scott, Sebastian L Johnston, Julie Rayes, Elizabeth Sapey, Aran Singanayagam, Juma El-Awaisi

## Abstract

Myocardial infarction (MI) triggers not only local cardiac damage but also a systemic inflammatory response that extends to remote organs. The pulmonary microcirculation, by virtue of its dense capillary network and direct anatomical proximity to the heart, is particularly vulnerable. Neutrophils and their effector mechanisms, including neutrophil extracellular traps (NETs) and the alarmin S100A8/A9, have been implicated in adverse cardiovascular outcomes. However, their role in remote damage post-MI remains unclear. Using intravital *in vivo* imaging in murine MI models and analysis of human lung tissues, we show that MI induces rapid pulmonary neutrophil and platelet recruitment, formation of platelet-neutrophil aggregates within capillaries, and endothelial activation. These changes are accompanied by NET release, fibrin deposition, and microvascular obstruction, leading to impaired vascular perfusion and necrosis. These pulmonary disturbances closely parallel those in the infarcted myocardium and exceed responses observed in other organs such as the kidney and liver, highlighting the lung as a vulnerable target organ. Increased neutrophil recruitment was associated with marked upregulation of the neutrophil-derived, NET-associated alarmin S100A8/A9 in mouse and human lungs, where it co-localised with infiltrating neutrophils, NETs, and platelet aggregates. Additionally, we show that short-term pharmacological inhibition of S100A8/A9 with ABR-238901 significantly attenuated pulmonary neutrophil infiltration, reduced NETosis and fibrin deposition, and restored capillary perfusion while rebalancing the pulmonary immune landscape. Together, these findings identify the lung as a principal site of remote thrombo-inflammatory injury after MI and implicate S100A8/A9, a neutrophil-derived, NET-associated alarmin, as a mechanistic driver of pulmonary microvascular dysfunction. We propose that targeting this pathway could provide dual protection for both cardiac and pulmonary microcirculations in the acute phase of myocardial injury.

**Graphical Abstract:** 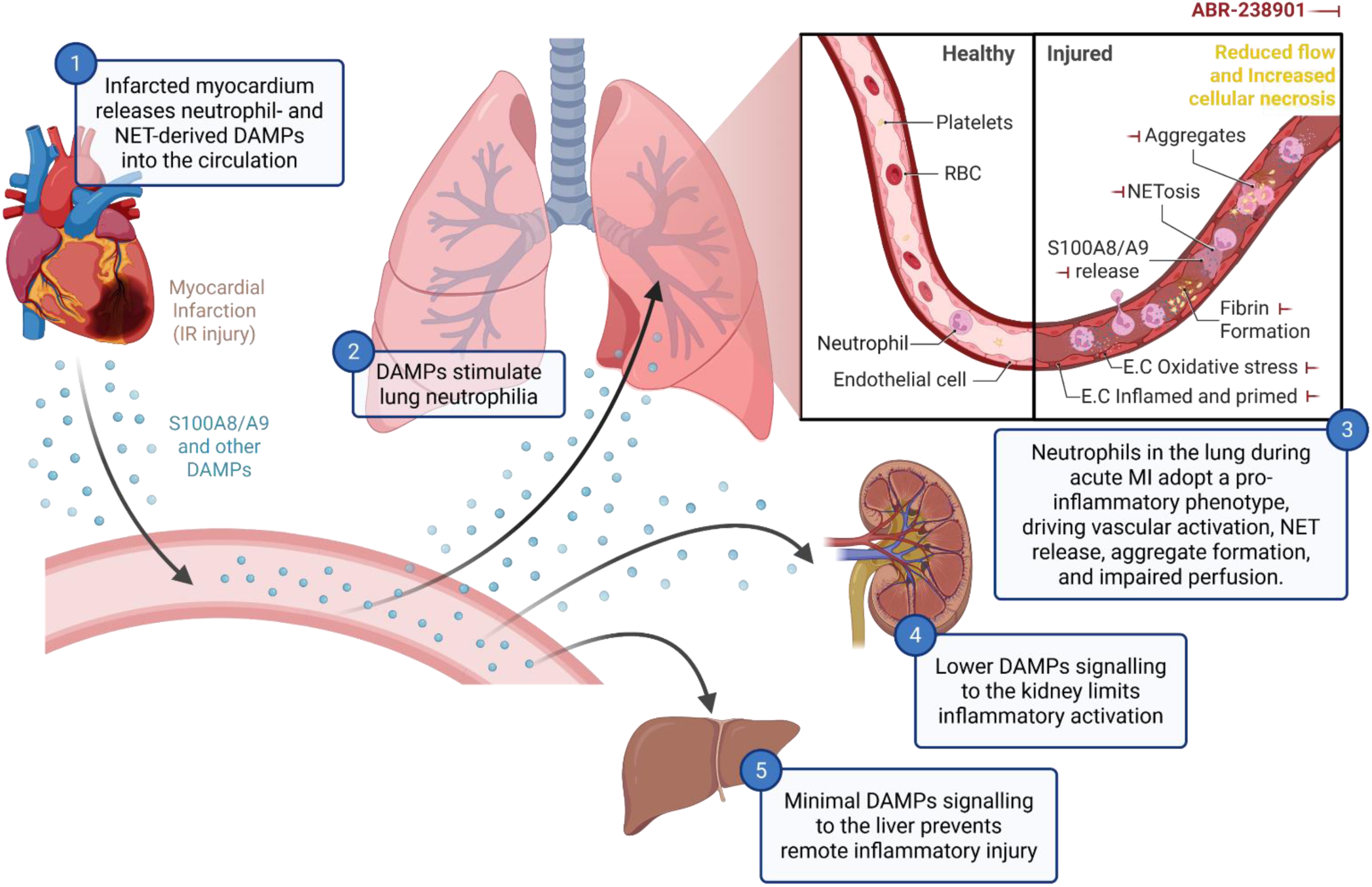

## Introduction

Myocardial infarction (MI) triggers not only local cardiac injury but also a profound systemic inflammatory response. In addition to leukocyte recruitment and cytokine release within the infarcted myocardium, remote organs can be adversely affected by this post-MI inflammatory surge [1]. MI has been associated with multi-organ dysfunction, suggesting that inflammatory pathways activated by myocardial ischaemia may extend to critical organs such as the lungs and kidneys. The lungs and kidneys are particularly susceptible to microvascular injury due to their dense capillary networks, high metabolic demands, and unique immunological environments, which render them vulnerable to systemic inflammatory and thrombotic insults [2, 3]. Clinically, MI can trigger acute lung injury (ALI) or acute respiratory distress syndrome (ARDS) even in the absence of other aetiologies; these conditions have been reported post-reperfusion, typically presenting with increased pulmonary vascular permeability and oedema [4, 5]. Similarly, acute kidney injury (AKI) occurs in approximately 20% of MI hospitalisations, and even mild renal impairment is associated with increased mortality [6]. These patterns highlight the potential for multi-organ involvement, manifesting as cardiopulmonary and cardiorenal syndromes, thereby complicating MI. This likely reflects the specific vulnerability of the pulmonary and renal microcirculations to systemic inflammation and circulatory disturbances triggered by myocardial ischaemia-reperfusion injury (IRI) following percutaneous coronary intervention (PCI). Although coronary microvascular disturbances post-MI are well characterised [7, 8], the pulmonary and renal microcirculatory responses remain poorly understood, partly due to the current limitations of clinical imaging in resolving dynamic microvascular changes.

The mechanistic links between myocardial injury and remote organ dysfunction, particularly pulmonary injury, remain poorly understood. However, a central driver of post-MI inflammation is the influx of neutrophils and monocytes into the infarcted myocardium. While essential for clearing necrotic debris, these myeloid cells simultaneously release a cascade of inflammatory mediators that disseminate systemically [8, 9]. Neutrophils, in particular, are key drivers of both local and systemic inflammation. Beyond their conventional antimicrobial functions, activated neutrophils can release neutrophil extracellular traps (NETs) through a process termed NETosis, which may occur via lytic cell death or a rapid, controlled ‘vital’ process with distinct molecular features [10]. These web-like structures, composed of DNA, histones, and neutrophil-derived proteins such as myeloperoxidase (MPO), neutrophil elastase, and the calcium-binding alarmin S100A8/A9 (also known as calprotectin), are increasingly recognised as contributors to post-ischaemic injury and thrombo-inflammation [11, 12]. Elevated levels of circulating NET-associated components, such as cell-free DNA, MPO-DNA complexes, neutrophil elastase, and S100A8/A9, have all been reported following MI and correlate with adverse outcomes [13-15]. NETs promote endothelial and platelet activation, increase vascular permeability, and serve as scaffolds for coagulation, thereby linking innate immune responses to thrombosis and microvascular dysfunction. Emerging evidence suggests NETs may also mediate remote organ damage by amplifying inflammatory and thrombotic cascades, in part through trapping damage-associated molecular patterns (DAMPs) such as HMGB1 and S100 proteins released from injured tissues [14, 16]. Among these, S100A8/A9 is particularly noteworthy, as elevated circulating levels after MI independently predict heart failure and adverse cardiovascular events [14, 17].

Building on evidence that neutrophils contribute to post-MI inflammation and may exert effects beyond the heart through the release of NETs [18], we investigated whether MI induces neutrophil- and NET-associated remote organ injury, focusing on the pulmonary microcirculation as a representative, particularly susceptible site. Specifically, we examined whether the neutrophil-derived alarmin S100A8/A9, which is abundantly expressed in neutrophils and becomes trapped in NETs, plays a mechanistic role in mediating post-MI pulmonary microvascular disturbances. Using intravital imaging, we visualised these processes to assess MI-induced pulmonary microvascular and perfusion disturbances. To contextualise these findings, we compared pulmonary responses with those in the kidney and liver and included chronic obstructive pulmonary disease (COPD) as a disease control to benchmark neutrophil-rich pulmonary inflammation.

## Methods

The data supporting the findings of this study are available from the corresponding author upon reasonable request.

### Ethical Approval

Ethical approval was obtained for the use of lung tissue and blood samples from patients undergoing routine thoracic surgery (17/WM/0272), as well as for the use of baseline blood samples from COPD patients (00/BA/459E) [19], with informed consent and institutional approvals. All animal experiments were reviewed and approved by the institutional Animal Welfare and Ethical Review Body (AWERB) and carried out under the authority of a Home Office Project Licence in compliance with the Animals (Scientific Procedures) Act of 1986.

#### Human Lung Tissue Collection

Human lung resections were obtained from patients undergoing thoracic surgery (predominantly lobectomy for lung cancer) at Queen Elizabeth Hospital Birmingham, UK, between January 2024 and September 2025. Patients were categorised into three groups: (i) those with confirmed MI within the preceding eight months, (ii) those with COPD, defined by the Global Initiative for Chronic Obstructive Lung Disease (GOLD) stage II-IV criteria, and (iii) individuals with no known respiratory or cardiovascular disease (“healthy” controls). Surplus tissue not required for histopathological analysis was collected. A segment distant from any tumour (if present) and without macroscopic pathology was immediately processed for histology or flow cytometry.

#### Mouse Models of MI and COPD

Myocardial IRI was induced in male BALB/c mice (8-10 weeks old) as previously described to model MI [7, 8, 20]. Anaesthesia was initiated via intraperitoneal injection of ketamine hydrochloride (100 mg/kg) and medetomidine hydrochloride (10 mg/kg). Mice were intubated and ventilated with oxygen using a MiniVent ventilator (Harvard Apparatus; stroke volume 220 μL; respiratory rate 130 breaths/min). A left thoracotomy was performed to expose the heart, and the left anterior descending (LAD) coronary artery was ligated for 45 minutes, followed by reperfusion lasting up to 2 hours. ABR-238901 (30 mg/kg, S100A8/A9 inhibitor) or PBS was delivered 20 minutes prior to reperfusion and again at 60 minutes post-reperfusion [17].

To model COPD, mice were lightly anaesthetised with isoflurane and administered 1.2 units of porcine pancreatic elastase (Merck) in 50 µL intranasally, as previously reported [21]. This induces histological emphysema and airway remodelling. After 10 days, mice were infected intranasally with 2.5 × 10⁶ tissue culture infectious dose 50 (TCID₅₀) of rhinovirus (RV) serotype A1B (A.T.C.C.), propagated in Ohio HeLa cells [21, 22]. This combined elastase-RV model recapitulates key features of COPD exacerbations, displaying augmented neutrophilic and pro-inflammatory responses. Tissues and blood were collected 24 hours post-RV infection for histological, cytometric, and molecular analysis.

#### Intravital Microscopy and Laser Speckle Imaging

Real-time intravital microscopy of the left lung was performed as previously described [8, 23]. A modified thoracic window (steel stabiliser) applied gentle negative pressure (∼20 mmHg) to a surface region of the left lung, stabilising it while maintaining ventilation. Intravenous injection of Ly-6G and CD41 antibodies (BioLegend) enabled simultaneous imaging of neutrophils and platelets. FITC-BSA (Sigma) was used to visualise vascular perfusion and functional capillary density (FCD). To image NETs, a triple-labelling strategy visualised extracellular DNA (SYTOX™ Green, Thermo Fisher), histones (phospho-H2A.X antibody, BioLegend), and neutrophil elastase (NE, Santa Cruz). Fibrin was visualised using fluorescently labelled fibrinogen (Thermo Fisher). Propidium iodide (PI, Thermo Fisher) was used to detect necrotic cells. Imaging was performed on an Olympus BX61WI spinning disk confocal microscope, and images were analysed with Slidebook 6 software [7, 8, 20].

Laser speckle contrast imaging (LSCI) quantified pulmonary blood flow using the moorFLPI-2 system (Moor Instruments, UK) as previously described [24, 25]. Flux data were captured across ischaemia and reperfusion phases and analysed using mFLPI2Measure and SpAn software.

### Immunofluorescence and Flow Cytometry

**- *Human lung tissue:*** Paraffin-embedded lung sections (6 μm) were processed and stained as previously described [26]. Briefly, sections were fixed in 4% paraformaldehyde (PFA), dehydrated, and stained with antibodies against S100A8/A9 (R&D Systems), CD66b (Biolegned), MPO (Abcam), and counterstained with Hoechst 33342 (Sigma-Aldrich), or with H&E (Abcam). Lung autofluorescence was quenched using a commercial kit (Vector Laboratories), and slides were mounted with ProLong™ Gold Antifade Mountant (Life Technologies). Imaging was performed using a Zeiss Axioscan Z1, and images were analysed using ZEN (Zeiss) and ImageJ software. A separate portion of lung tissue was digested in RPMI containing 0.1% collagenase D (Sigma-Aldrich) and 0.01% DNase I (Roche) to obtain single-cell suspensions [27]. Cells were stained with fluorophore-conjugated antibodies against oxidative damage (8-OHdG, Abcam) and inflammatory mediators including S100A9, IL-6, IL-1β, TNF, and IFN-γ. A minimum of 250,000 events per sample were acquired using a BD LSR Fortessa™ III cytometer and analysed using FlowJo v10.9 (BD Biosciences).
**- *Mouse tissue:*** Frozen sections (10 μm) from multiple organs were fixed in acetone and stained with antibodies targeting S100A9 (MRP-14, BD Biosciences), neutrophils (Ly6G, BioLegend), platelets (CD41, BioLegend), E-selectin (CD62E, Santa Cruz), VCAM-1 (CD106, BioLegend), and oxidative DNA/RNA damage (8-OHdG, Abcam) [7]. Images were captured on an EVOS FL microscope, and mean fluorescence intensity (MFI) was quantified using ImageJ. Fresh lungs were digested to single-cell suspensions using Liberase and DNase (Roche) in RPMI supplemented with 10% FBS, filtered, lysed, and stained with LIVE/DEAD™ viability dye (Invitrogen) [22]. Cells were incubated with CD45 and Fc Block (BD Biosciences), followed by staining with antibody panels targeting: (i) CD11c (BD Pharmingen), CD11b, Siglec-F (BD Pharmingen), Ly6G, Ly6C, F4/80, CD63, and CD64; (ii) CD4, CD8a, CD19, CD3e (Thermo Fisher), TCR γ/δ, and CD335 (NKp46); (iii) CD31, CD326 (EpCAM), oxidative DNA/RNA damage (anti-8-OHdG, Abcam), CD106/VCAM-1, E-selectin (CD62E, Santa Cruz), and CD41. Samples were fixed with 1% PFA before acquisition on a BD LSR Fortessa™ III cytometer and analysed using FlowJo v10.9. All antibodies were sourced from BioLegend unless otherwise stated.

### Luminex Cytokine Assays

**- *Human Plasma*:** Plasma samples were isolated from EDTA-anticoagulated blood collected either pre-operatively (from the patient groups described above) or from baseline GOLD stage II COPD patients recruited at Imperial College Healthcare NHS Trust, London, UK. A custom multiplex assay (R&D Systems Human Luminex Discovery Assay) was used to quantify S100A9, IL-6, IL-1β, TNF, and IFN-γ on a Luminex® 200 platform.
**- *Mouse Plasma:*** Cytokine concentrations were measured using the Bio-Plex Pro™ Mouse Cytokine 23-plex Assay (Bio-Rad) on a Luminex® 200. Comparisons were made across sham-operated, IRI, IRI + ABR-238901-treated, and experimental COPD groups.

#### Statistical analysis

Statistical analyses were performed using GraphPad Prism 9 (GraphPad Software Inc.). One-way ANOVA with Tukey’s post hoc test was used for comparisons across three or more groups. Data are presented as mean ± SEM; p < 0.05 was considered statistically significant. LSCI results are reported as perfusion flux units normalised to baseline readings.

## Results

### Myocardial IRI triggers neutrophil- and NET-driven pulmonary thrombo-inflammation

Building on evidence that MI can induce inflammatory responses beyond the heart, we first assessed whether myocardial IRI leads to neutrophil- and NET-associated thrombo-inflammatory disturbances within the pulmonary microcirculation. Intravital imaging revealed minimal neutrophil and platelet recruitment in sham lungs, with no microvascular aggregates or microthrombi in both surface and deeper tissue layers.

Following myocardial IRI, pulmonary vessels displayed a marked increase in neutrophil accumulation and significant platelet adhesion, with frequent co-localisation within the capillary network **(Figure 1A-B, F-I)**. Notably, this contrasted with sham mice, which also underwent thoracotomy but showed minimal pulmonary neutrophil or platelet recruitment, suggesting that myocardial IRI itself, rather than surgical trauma, drives the exaggerated pulmonary response. These IRI-induced interactions culminated in the formation of platelet-neutrophil aggregates and intravascular microthrombi, suggesting coordinated cellular crosstalk within the lung microcirculation **(Figure 1A-B)**. To examine whether this cellular aggregation was associated with downstream thrombus maturation, we next assessed fibrin formation. Fibrin deposition was substantially elevated following MI but essentially absent in sham lungs, confirming the presence of pulmonary microthrombi **(Figure 1C, J)**.

**Figure 1.**
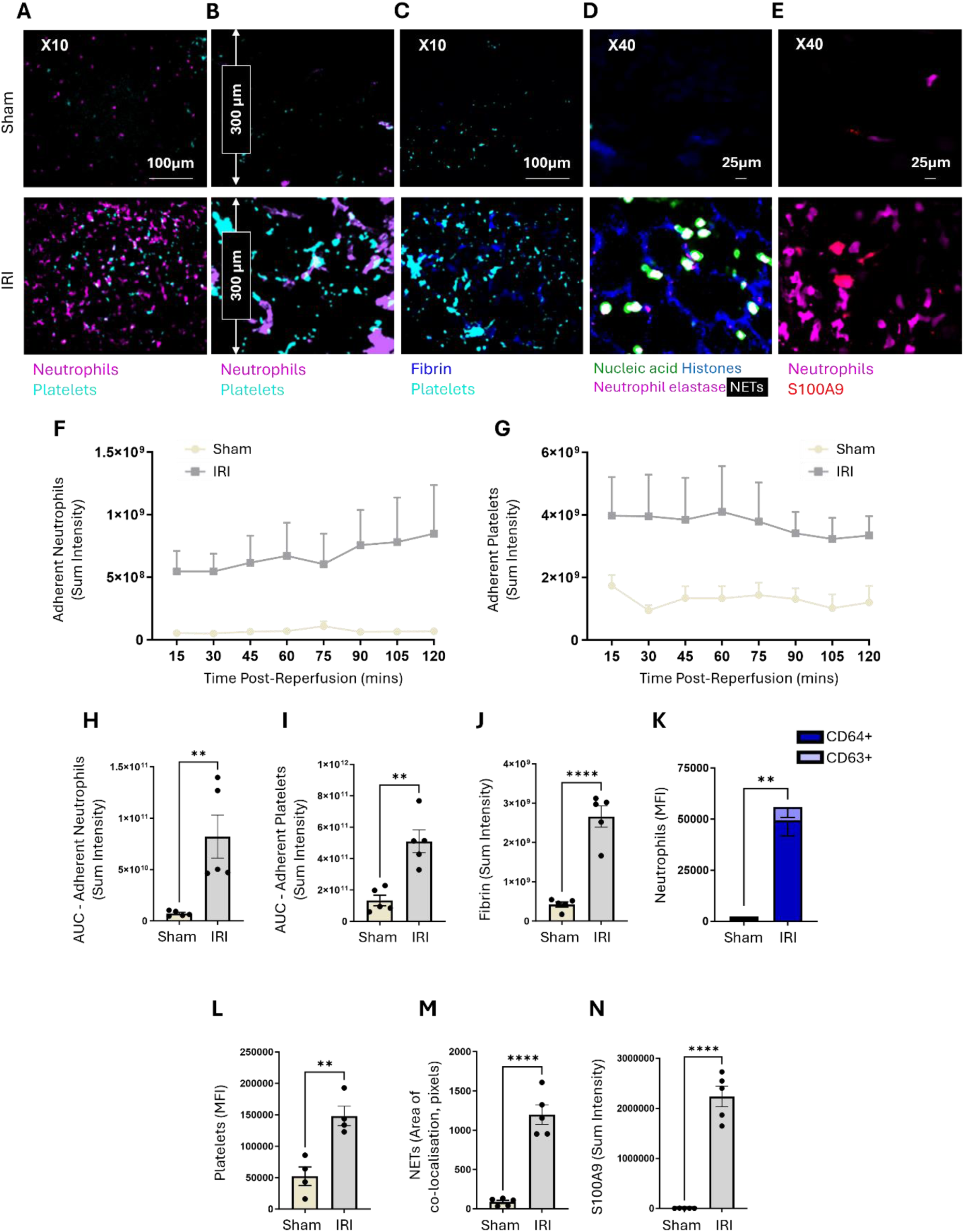
Myocardial IRI triggers neutrophil- and NET-driven pulmonary thrombo-inflammation *in vivo*. Mice underwent myocardial ischaemia-reperfusion injury (IRI) to model myocardial infarction. **(A)** Representative intravital images showing adherent neutrophils and platelets in the pulmonary microcirculation. **(B)** Representative multiphoton Z-stack images at a depth of 300 μm. Representative intravital images showing **(C)** fibrin formation, **(D)** NETs, and **(E)** S100A8/A9 release at 120 minutes post-reperfusion. Quantitative analysis of intravital data for **(F)** adherent neutrophils and **(G)** platelets, and area under the curve (AUC) analysis for **(H)** adherent neutrophils and **(I)** platelets in the pulmonary microcirculation over a 120-minute time course. N = 5/group. **(J)** Quantitative analysis for fibrin formation at 120 minutes post-reperfusion. Quantitative flow cytometric analysis of **(K)** neutrophils and **(L)** platelets in collagenase-digested lungs. N = 4/group. Quantitative analysis of intravital data for **(M)** NETs and **(N)** S100A9 release at 120 minutes post-reperfusion. N = 5/group. *p < 0.05, **p < 0.01, ***p < 0.001, ****p < 0.0001 when tested using a one-way ANOVA followed by Tukey multiple comparison test.

To further characterise neutrophil activation and the associated thrombo-inflammatory milieu, flow cytometric analysis of digested pulmonary tissue showed that MI significantly increased the mean fluorescence intensity (MFI) of neutrophil markers, particularly CD64, consistent with enhanced neutrophil activation **(Figure 1K)**. Platelet-associated signal was also significantly elevated post-MI **(Figure 1L)**, reflecting a heightened microthrombotic burden.

To assess neutrophil effector activity, we next visualised NET formation *in vivo.* NET formation was minimal in sham lungs but dramatically increased post-MI, as evidenced by a significant rise in markers of NET formation, including histone H2A.X and extracellular DNA, and their co-localisation with neutrophil elastase, indicating enhanced NET release following MI **(Figure 1D, M)**. S100A8/A9 was abundantly detected within recruited neutrophils in the pulmonary vasculature. Importantly, extracellular S100A8/A9 signal was observed in areas of dense immune cell infiltration, consistent with its retention within NET structures, suggesting robust neutrophil activation and NETosis **(Figure 1E, N)**. Together, these findings demonstrate that MI induction in mice leads to robust neutrophil- and platelet-driven pulmonary microvascular inflammation, coupled with NET formation.

### Myocardial IRI triggers pulmonary perfusion failure and necrosis

Neutrophil adhesion, NETosis, platelet aggregation, and fibrin deposition can all impede capillary blood flow [28]. We therefore next assessed the functional consequences of MI on pulmonary perfusion. Intravital microscopy of FITC-BSA-labelled vessels revealed a dense and continuous network of perfused pulmonary capillaries in sham mice, particularly evident around alveolar spaces **(Figure 2A)**. In the immediate aftermath of MI, lungs displayed extensive areas devoid of FITC-BSA signal, corresponding to a lack of perfusion. This likely reflects not only microvascular obstruction but also circulatory collapse secondary to acute myocardial dysfunction. These perfusion deficits often overlapped with zones of PI⁺ necrotic cells, suggesting that capillary obstruction may drive localised cell death **(Figure 2A-B)**. Additionally, overall pulmonary necrosis was significantly increased post-MI **(Figure 2C)**.

**Figure 2.**
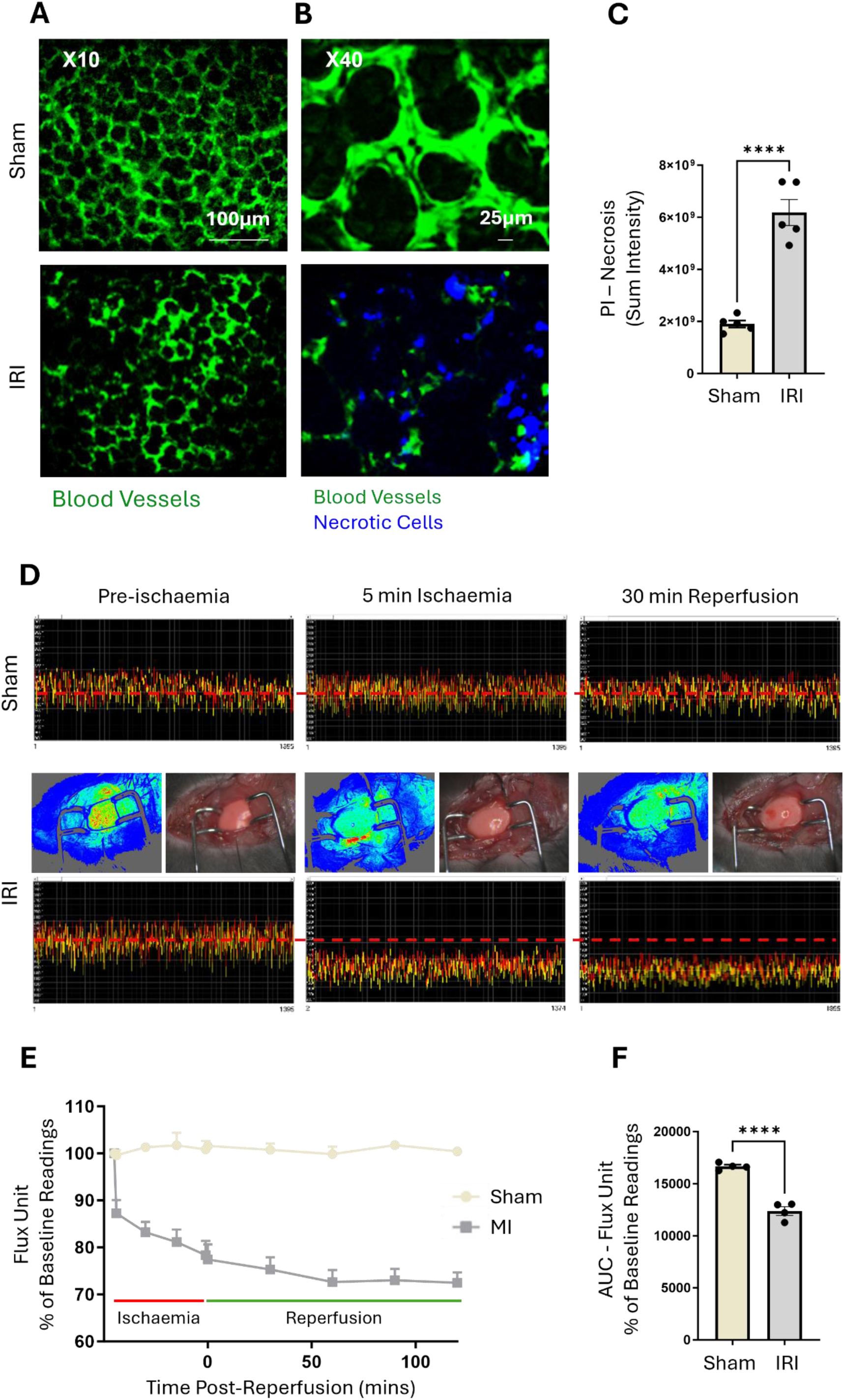
Myocardial IRI triggers pulmonary perfusion failure and necrosis *in vivo*. Mice underwent myocardial ischaemia-reperfusion injury (IRI) to model myocardial infarction. Representative intravital images showing **(A)** FITC-BSA perfused pulmonary microvessels and **(B)** cellular necrosis identified by propidium iodide labelling at 120 min post-reperfusion. **(C)** Quantitative analysis of intravital data for cellular necrosis. N = 5/group. **(D)** Representative LSCI heatmaps and corresponding 1-min real-time flux readouts of the ventilated lung. Heatmaps show warm colours under basal conditions, cooler colours during ischaemia, and warmer colours again as reperfusion is initiated. **(E)** Quantitative time-course analysis of flux unit readings as a percentage of baseline values during the reperfusion phase and **(F)** corresponding area under the curve (AUC). N = 4/group. *p < 0.05, **p < 0.01, ***p < 0.001, ****p < 0.0001 when tested using a one-way ANOVA followed by Tukey multiple comparison test.

To complement these findings and quantify perfusion across a broader pulmonary surface, we used LSCI. Flux values remained stable throughout the reperfusion phase in sham mice, with consistent warm colours on perfusion heatmaps indicating good blood flow. In contrast, MI induced a rapid and pronounced decline in perfusion, reflected by significantly decreased flux values that failed to recover even after reperfusion **(Figure 2D-F)**. Together, these findings demonstrate that MI induces a potent thrombo-inflammatory response in the pulmonary microvasculature, characterised by neutrophil and platelet activation, endothelial injury, and vascular occlusion. These changes likely arise from a combination of systemic inflammatory signalling and acute circulatory collapse specific to cardiac injury, rather than from surgical trauma alone.

### Myocardial IRI disrupts the pulmonary immune microenvironment

To characterise the broader immunological changes linking cardiac injury to lung responses, we next analysed the immune cell profile in digested pulmonary tissue to identify subsets driving the microvascular disturbances described above. MI was associated with increased MFI of monocyte markers, with higher expression observed in both classical Ly6C^high^ and non-classical Ly6C^low^ monocyte populations **(Figure 3A)**, alongside an increase in CD45-associated total leukocyte signal **(Figure 3B)**. Gamma delta (γδ) T-cell signal was also significantly increased post-MI, suggesting activation of innate-like lymphocytes **(Figure 3C)**, while NK cells showed a non-significant upward trend **(Figure 3D)**. Notably, alveolar macrophages **(Figure 3E)**, eosinophils, CD4^+^ and CD8^+^ T cells, and CD19^+^ B cells showed no substantial change **(Supplementary Figure 1)**. This suggests that while certain innate cell types (such as neutrophils and monocytes) were activated, other innate populations and the adaptive immune response remained largely unaffected.

**Figure 3.**
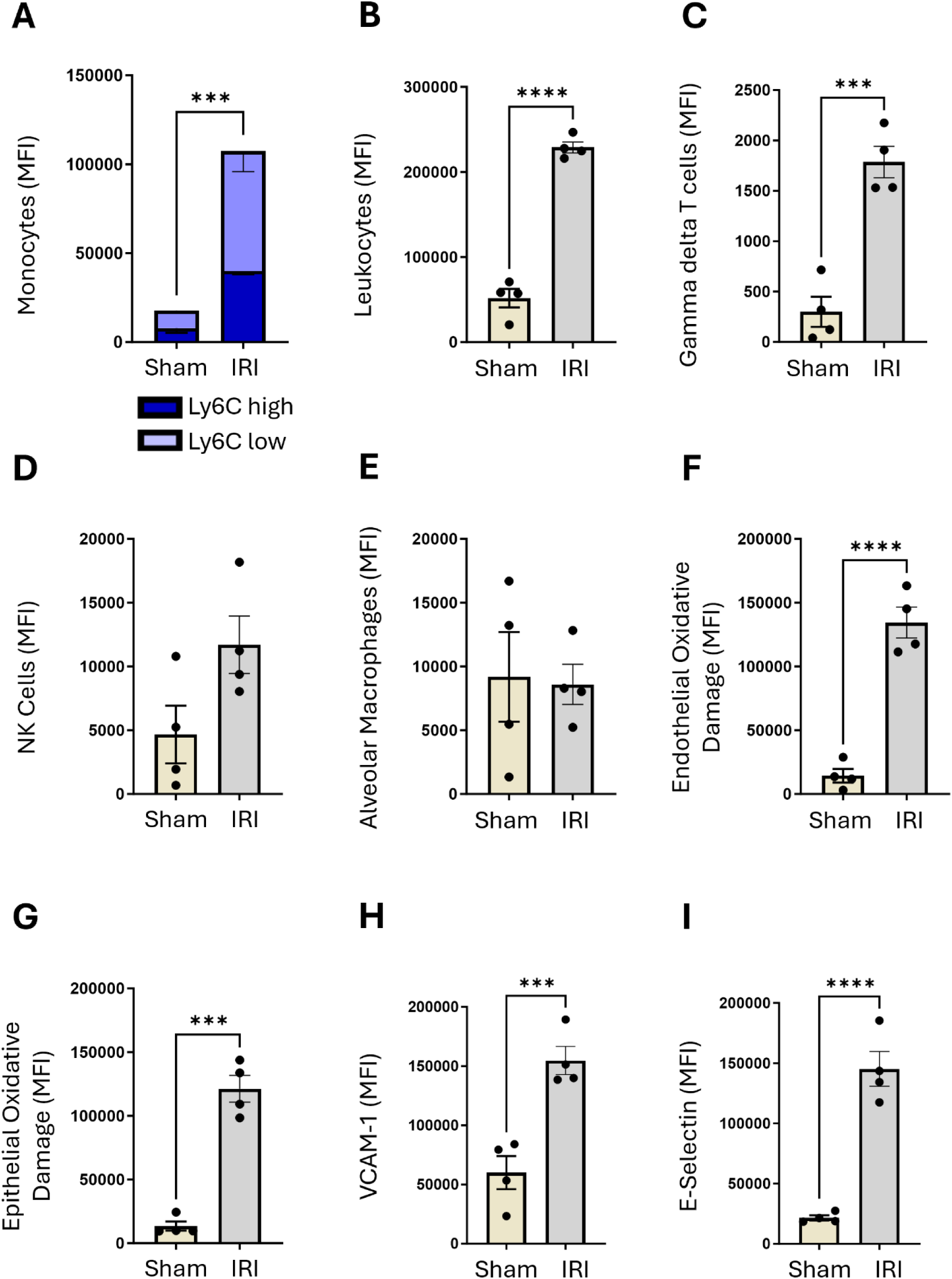
Myocardial IRI disrupts the pulmonary immune microenvironment. Mice underwent myocardial ischaemia-reperfusion injury (IRI) to model myocardial infarction, and lungs were harvested and collagenase-digested. Quantitative flow cytometric analysis of **(A)** monocytes, **(B)** total leukocytes, **(C)** gamma delta T cells, **(D)** NK cells, **(E)** alveolar macrophages, **(F)** endothelial oxidative damage, **(G)** epithelial oxidative damage, **(H)** VCAM-1, and **(I)** E-selectin. N = 4/group. *p < 0.05, **p < 0.01, ***p < 0.001, ****p < 0.0001 when tested using a one-way ANOVA followed by Tukey multiple comparison test.

To assess changes in the endothelial and epithelial microenvironment, we measured markers of oxidative stress and adhesion. MI increased 8-OHdG expression in both endothelial and epithelial compartments, indicating widespread oxidative DNA damage **(Figure 3F-G)**. Endothelial activation was further confirmed by elevated VCAM-1 and E-selectin expression **(Figure 3H-I)**, consistent with a primed vasculature favouring leukocyte recruitment. Together, these findings reveal that MI triggers a focused but profound remodelling of the pulmonary immune microenvironment, primarily driven by neutrophils and monocytes.

### MI elicits pulmonary and systemic inflammation comparable to COPD

Inflammation is a hallmark of both MI and COPD, yet it remains unclear whether the MI-induced pulmonary damage described above mirrors that observed in a primary lung condition characterised by neutrophil-rich inflammation, such as COPD. To investigate this, we compared pulmonary inflammation in a myocardial IRI model with that in a well-established model of exacerbated COPD, in which elastase-induced emphysema is combined with rhinovirus infection to trigger pro-inflammatory responses.

Immunostaining revealed comparable increases in pulmonary neutrophil recruitment and platelet adhesion between MI and COPD mice **(Figure 4A-C)**. Likewise, expression of endothelial adhesion molecules VCAM-1 and E-selectin did not differ significantly between the two models **(Figure 4D-E)**. Oxidative damage, assessed by 8-OHdG staining, was also similar in both models **(Figure 4F)**.

**Figure 4.**
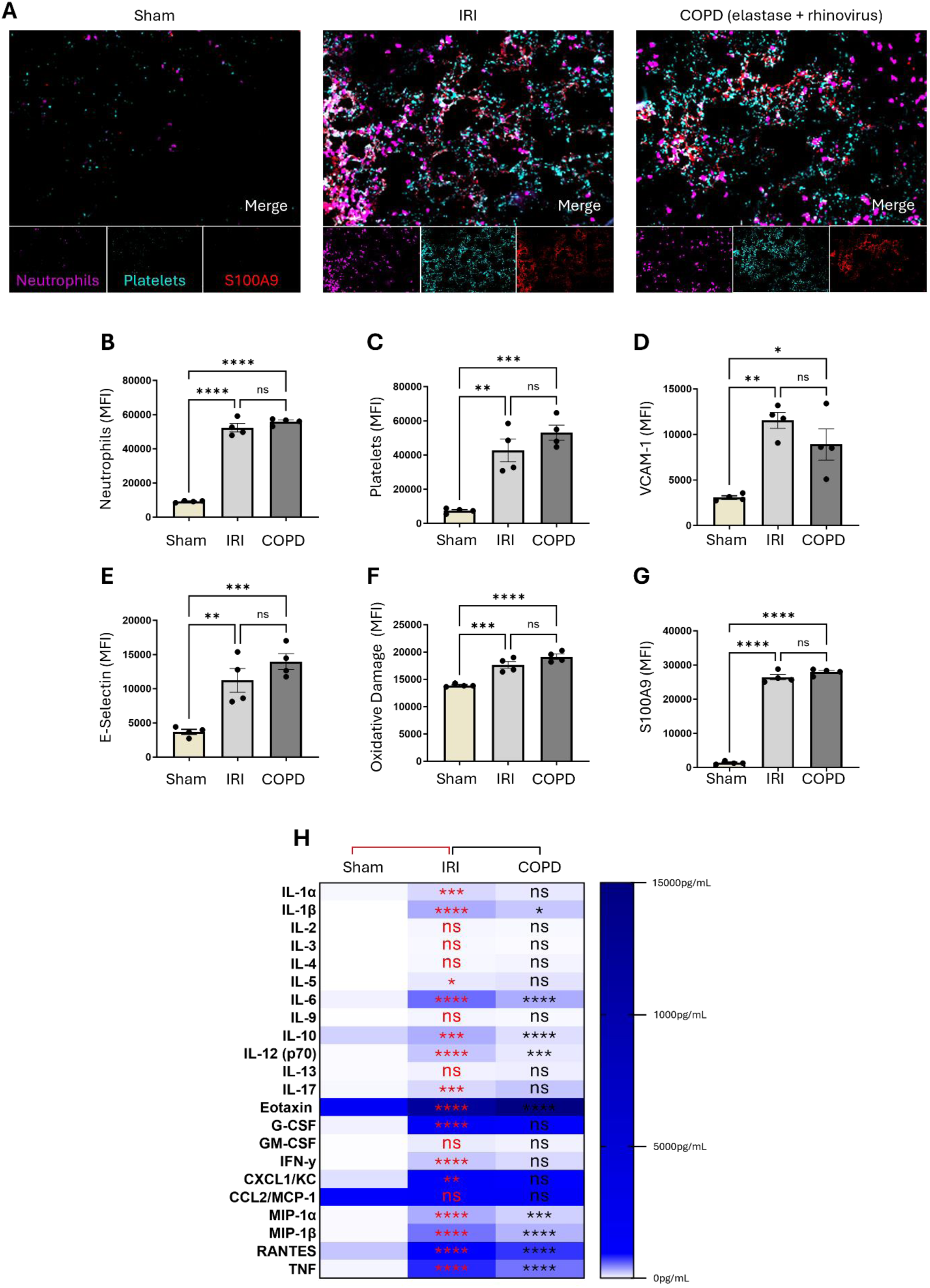
Myocardial IRI upregulates neutrophil-derived and NET-associated S100A8/A9 expression to levels comparable to COPD. Mice underwent myocardial ischaemia-reperfusion injury (IRI) to model myocardial infarction or were administered elastase and rhinovirus intranasally to model chronic obstructive pulmonary disease (COPD). **(A)** Representative immunofluorescence images of frozen lung sections showing neutrophils, platelets, and S100A9. Quantitative analysis of immunofluorescence images for **(B)** neutrophil recruitment, **(C)** platelet adhesion, **(D)** VCAM-1, **(E)** E-selectin, **(F)** oxidative damage, and **(G)** S100A9 expression. **(H)** Quantitative analysis of multiplex cytokine profiles for inflammatory markers. Statistical comparisons are shown above the table. Results are presented as mean ± SEM pg/ml. N = 4/group. *p < 0.05, **p < 0.01, ***p < 0.001, ****p < 0.0001 when tested using a one-way ANOVA followed by Tukey multiple comparison test.

Having established that MI induces pulmonary inflammation at levels comparable to COPD, we next asked whether this extended to the systemic inflammatory milieu. Systemic inflammation is a defining feature of MI, correlating with infarct size and clinical outcomes, and it is implicated in remote organ injury. We therefore profiled 23 inflammatory mediators in the plasma of MI mice using a multiplex Luminex assay. Compared with sham controls, myocardial IRI significantly increased circulating levels of 15 cytokines, including IL-1β, IL-6, TNF, IFN-γ, monocyte chemoattractant protein-1 (MCP-1), and granulocyte-macrophage colony-stimulating factor (GM-CSF), among others **(Figure 4H)**. These increases are consistent with a systemic inflammatory response and align with our previous data [8]. To contextualise the degree of systemic inflammation in MI, we compared cytokine levels with those from elastase and rhinovirus COPD mice. MI was associated with higher levels of eight cytokines than in COPD, with only eotaxin (CCL11), a chemokine involved in eosinophil recruitment, found to be significantly lower in MI **(Figure 4H)**. Taken together, these findings indicate that while both MI and COPD drive substantial pulmonary and systemic inflammation, the nature of the response differs: MI is dominated by high levels of key pro-inflammatory cytokines such as IL-1β and IFN-γ, consistent with an acute sterile injury.

### MI upregulates neutrophil-derived and NET-associated S100A8/A9 expression to levels comparable to COPD

Both clinical and experimental data implicate the neutrophil-derived and NET-associated alarmin S100A8/A9 in post-MI pathological processes. Recent evidence has associated elevated circulating levels of S100A8/A9 with increased risk of adverse cardiovascular outcomes and has shown that plasma S100A8/A9 independently predicts the development of post-MI heart failure [14, 17]. We therefore next assessed whether myocardial IRI elevates S100A8/A9 levels in the lung. Immunostaining showed low constitutive S100A9 expression in sham lungs, which increased significantly following MI, with a distribution pattern and levels similar to those seen in COPD **(Figure 4A, G)**. S100A9 co-localised strongly with Ly6G^+^ neutrophils and, to a lesser extent, CD41^+^ platelets in the capillary-rich pulmonary microvasculature **(Figure 4A)**.

To validate these findings in human tissue, we examined lung samples obtained from patients undergoing thoracic surgery for other clinical indications (predominantly lung cancer), including patients with recent MI, patients with COPD, and individuals without respiratory or cardiovascular disease (“healthy”). Immunostaining revealed comparable increases in pulmonary neutrophil recruitment in MI and COPD patient groups **(Figure 5A-B)**. In “healthy” lung margins, S100A8/A9 expression was low but constitutive, whereas it was significantly increased in both MI and COPD lungs **(Figure 5A, D)**. S100A8/A9 localisation closely mirrored neutrophil infiltration, with notable co-localisation in areas of dense neutrophil accumulation **(Figure 5A, B)**. Flow cytometric analysis of lung tissue digests demonstrated a pattern of oxidative DNA damage (8-OHdG) that paralleled S100A8/A9 expression **(Figure 5C)**. Interestingly, S100A9 expression in the lung surpassed that of other pro-inflammatory cytokines, including IL-6, IL-1β, TNF, and IFN-γ, with no significant differences between MI and COPD groups **(Figure 5E)**. In parallel, plasma analysis was performed using blood from MI patients, individuals with GOLD stage II-IV COPD (from our surgical resection cohort), and an independent cohort of baseline GOLD stage II COPD patients. Circulating S100A9 levels were markedly elevated in both conditions compared with healthy controls, to a greater extent than other mediators, while total neutrophil counts in peripheral blood were unchanged across groups **(Figure 5F-H)**. These findings suggest that MI induces a pulmonary thrombo-inflammatory microenvironment in which S100A8/A9 serves as a key neutrophil- and NET-associated mediator of these disturbances.

**Figure 5.**
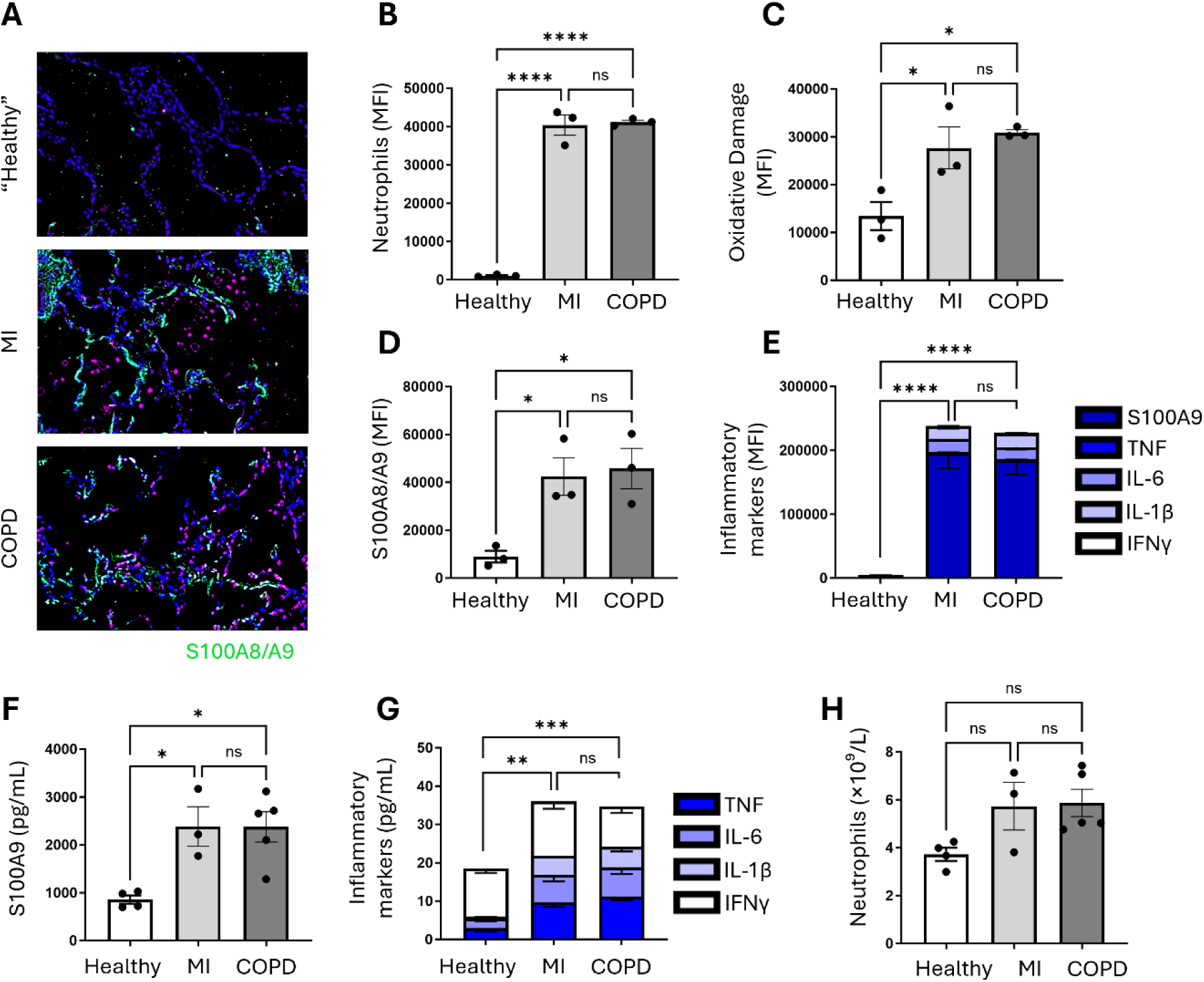
MI upregulates neutrophil-derived and NET-associated S100A8/A9 expression to levels comparable to COPD. **(A)** Representative immunofluorescence images of paraffin-embedded lung sections showing S100A9 and neutrophils from MI patients, individuals with GOLD stage II-IV COPD, and healthy controls. Quantitative analysis of immunofluorescence images for **(B)** neutrophil recruitment, **(C)** oxidative damage, **(D)** S100A9, and **(E)** inflammatory markers. N = 3/group. Quantitative analysis of Luminex cytokine data for **(F)** S100A9, **(G)** inflammatory markers, and **(H)** neutrophil counts in plasma from MI patients, individuals with GOLD stage II-IV COPD (surgical resection cohort), and individuals with GOLD stage II COPD. N ≥ 3/group. *p < 0.05, **p < 0.01, ***p < 0.001, ****p < 0.0001 when tested using a one-way ANOVA followed by Tukey multiple comparison test.

### The lung as a primary target of remote myocardial IRI

Systemic consequences of MI are known to extend beyond the heart, but direct comparisons of thrombo-inflammatory injury across multiple organs remain limited. Given the pulmonary circulation’s intimate proximity to the heart and its extensive capillary network, we sought to determine whether pulmonary microvascular responses to MI resemble those seen in the infarcted myocardium and how they compare to other remote organs **(Figure 6A-C)**. Mouse immunofluorescence analysis revealed extensive expression of S100A9, neutrophils, and platelets in the heart following MI **(Figure 6D-F)**, with comparable levels observed in the lungs **(Figure 4B-C, G)**. Notably, VCAM-1 and E-selectin expression were markedly upregulated in both the heart **(Figure 6G-H)** and lungs **(Figure 4D-E)**, with the lungs exhibiting levels comparable to the infarcted myocardium. Markers of endothelial injury demonstrated organ-specific variation: oxidative DNA damage was most prominent in the heart **(Figure 6I)** but remained significantly elevated in the lungs **(Figure 4F)**.

**Figure 6.**
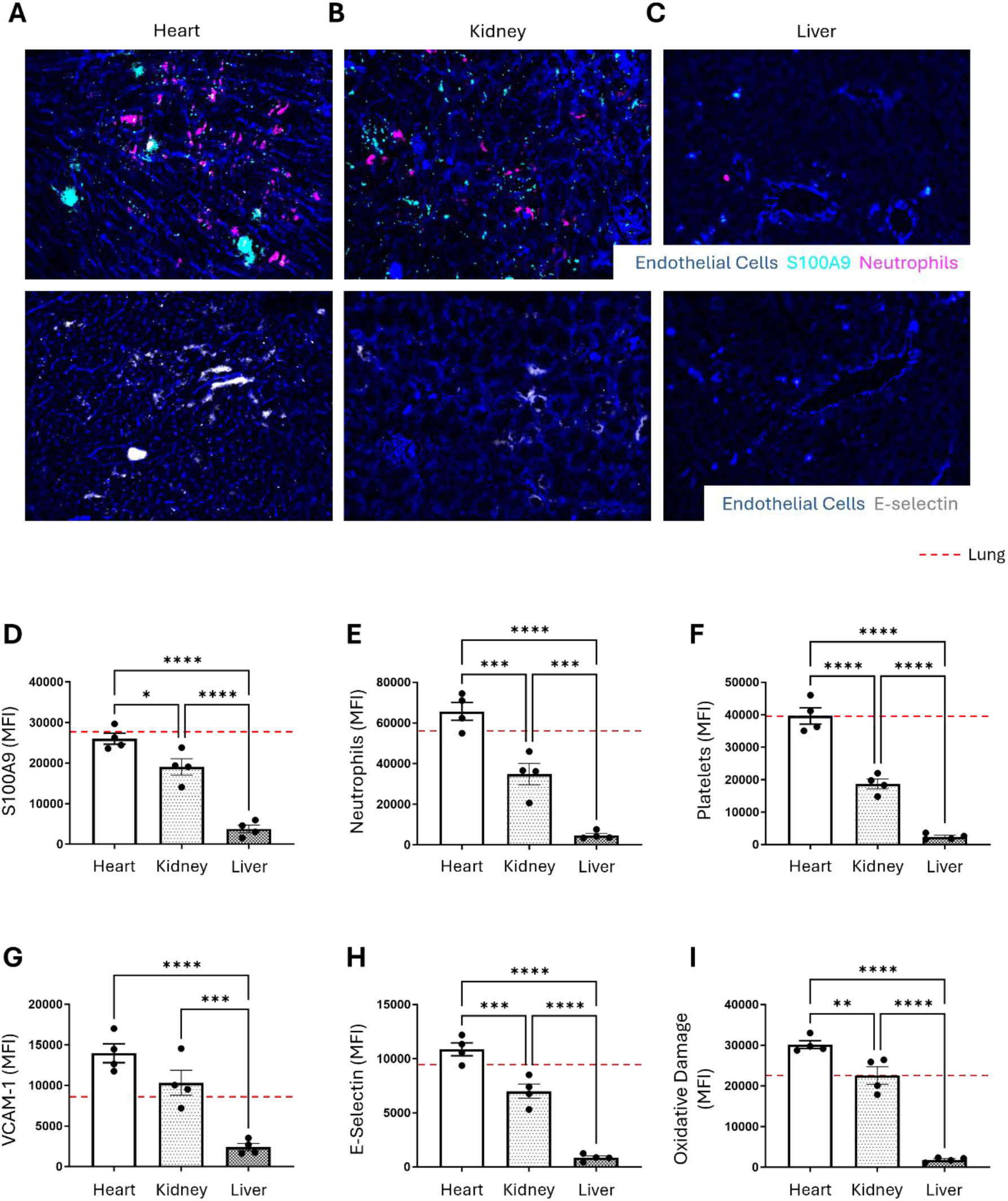
The lung as a primary target of remote myocardial IRI. Mice underwent myocardial ischaemia-reperfusion injury (IRI) to model myocardial infarction. Representative immunofluorescence images of frozen **(A)** heart, **(B)** kidney, and **(C)** liver sections showing S100A9, neutrophils, E-selectin, and endothelial cells. Quantitative analysis of immunofluorescence images for **(D)** S100A9, **(E)** neutrophil recruitment, **(F)** platelet adhesion, **(G)** VCAM-1, **(H)** E-selectin, and **(I)** oxidative damage expression. N = 4/group. *p < 0.05, **p < 0.01, ***p < 0.001, ****p < 0.0001 when tested using a one-way ANOVA followed by Tukey multiple comparison test.

While S100A9 was basally expressed at low levels across all sham groups **(Supplementary Figure 1)**, the kidneys showed lower yet notable increases in S100A9, neutrophils, platelets, E-selectin, and VCAM-1 compared with the lungs **(Figure 4)**, although they exhibited greater oxidative damage than the pulmonary tissue **(Figure 6B, D-I)**. The liver demonstrated the lowest levels of all markers, including negligible S100A9 and immune cell recruitment **(Figure 6C-I)**. These findings indicate that pulmonary thrombo-inflammatory responses following MI not only closely resemble those in the heart but also generally surpass those observed in other highly perfused organs within the two-hour timeframe examined.

### Inhibition of neutrophil-derived and NET-associated S100A8/A9 restores pulmonary immune and vascular homeostasis

Given the central role of S100A8/A9 in neutrophil activation, NET formation, and endothelial dysfunction [29], we investigated whether short-term pharmacological inhibition of S100A8/A9 using ABR-238901 could attenuate MI-induced pulmonary injury. Recent experimental studies have shown that ABR-238901 modulates post-myocardial IRI inflammation within the heart and circulation by suppressing haematopoietic stem cell proliferation in the bone marrow and limiting their mobilisation to the site of injury. This treatment has been shown to reduce infarct size, improve long-term cardiac function, and promote myocardial neovascularisation [17, 30, 31]. Here, we investigated its impact on the lung as a remote site of injury.

Treatment with ABR-238901 significantly attenuated pulmonary neutrophil recruitment **(Figure 7A-B, F)**, notably reducing the proportion of CD64⁺ pro-inflammatory neutrophils **(Figure 7G)**. It also decreased Ly6C^high^ monocytes, total leukocytes, and γδ T cells **(Figure 7H-J)**. Concurrently, anti-inflammatory CD63⁺ neutrophils and Ly6C^low^ monocytes were increased, suggesting a central role for S100A8/A9 in orchestrating neutrophil-driven pulmonary damage following MI. Platelet adhesion was also reduced **(Figure 7A-B, K)**. In parallel, platelet-neutrophil aggregates and microthrombi formation were markedly diminished, as observed using *in vivo* depth imaging **(Figure 7A-B)**, suggesting a rebalancing of the pulmonary immune-thrombotic landscape.

**Figure 7.**
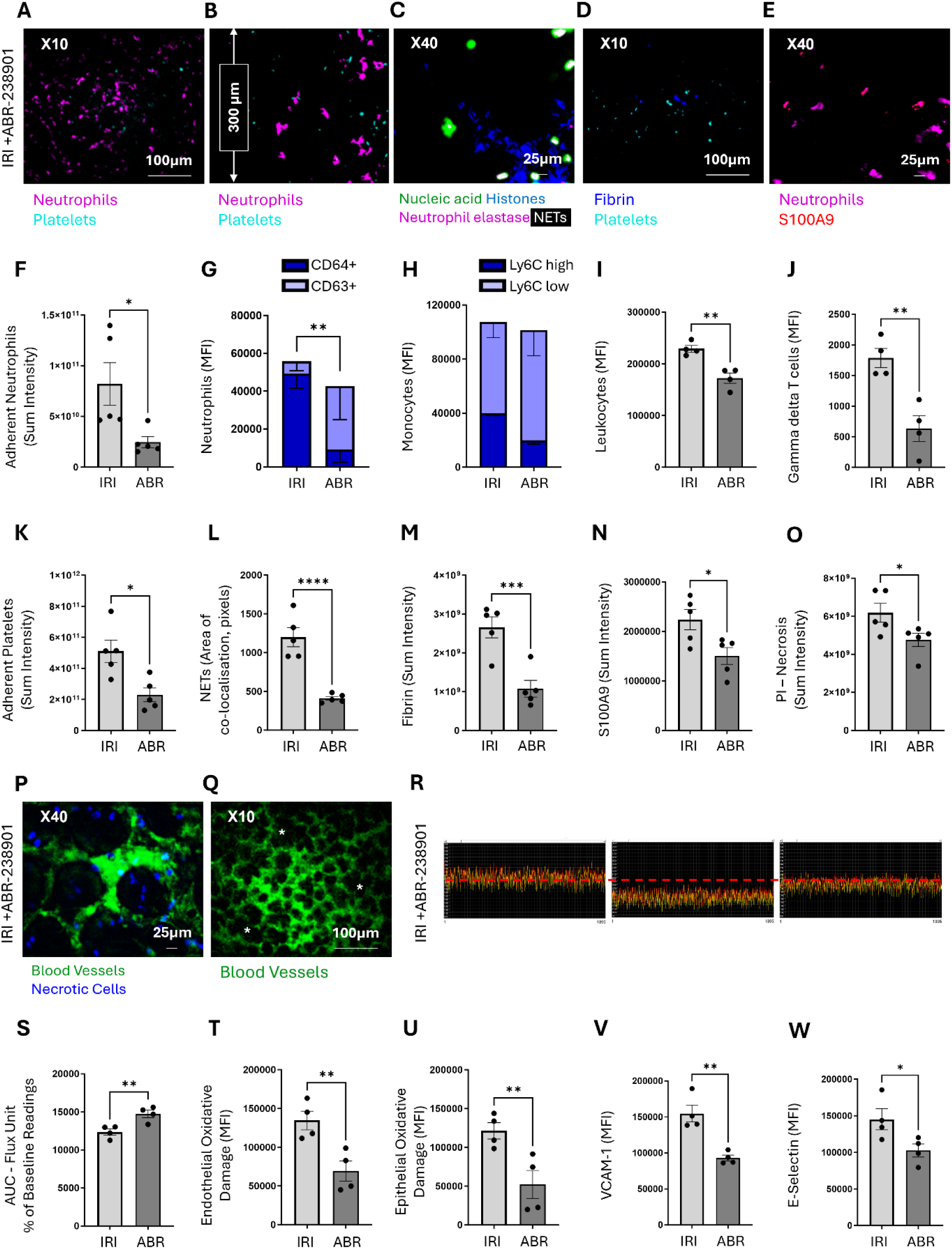
Inhibition of neutrophil-derived and NET-associated S100A8/A9 restores pulmonary immune and vascular homeostasis. Mice underwent myocardial ischaemia-reperfusion injury (IRI) to model myocardial infarction. ABR-238901 (S100A8/A9 inhibitor) was administered pre- and during reperfusion. **(A)** Representative intravital images showing adherent neutrophils and platelets in the pulmonary microcirculation. **(B)** Representative multiphoton Z-stack images at a depth of 300 μm. Representative intravital images showing **(C)** NETs, **(D)** fibrin formation, and **(E)** S100A8/A9 release at 120 minutes post-reperfusion. **(F)** Quantitative area under the curve (AUC) analysis for adherent neutrophils in the pulmonary microcirculation over a 120-minute time course. N = 5/group. Quantitative flow cytometric analysis of **(G)** neutrophils, **(H)** monocytes, **(I)** total leukocytes, and **(J)** gamma delta T cells in collagenase-digested lungs. N =45/group. Quantitative analysis of intravital data for **(K)** platelets, **(L)** NETs, **(M)** fibrin formation, **(N)** S100A9 release, and **(O)** cellular necrosis at 120 min post-reperfusion. N = 5/group. Representative intravital images showing **(P)** cellular necrosis identified by propidium iodide labelling and **(Q)** FITC-BSA perfused pulmonary microvessels at 120 min post-reperfusion. **(R)** Representative 1-min real-time flux readouts of the ventilated lung, and **(S)** corresponding time-course analysis of flux unit readings as a percentage of baseline values during the reperfusion phase only. N = 4/group. Quantitative flow cytometric analysis of **(T)** endothelial oxidative damage, **(U)** epithelial oxidative damage, **(V)** VCAM-1, and **(W)** E-selectin in collagenase-digested lungs. N = 4/group. *p < 0.05, **p < 0.01, ***p < 0.001, ****p < 0.0001 when tested using a one-way ANOVA followed by Tukey multiple comparison test.

ABR-238901 also mitigated neutrophil effector functions. NET formation, which was markedly elevated post-MI, was significantly reduced following the treatment, as indicated by decreased staining for histones, DNA, neutrophil elastase, and S100A8/A9 **(Figure 7C, L)**. Fibrin deposition and S100A8/A9 release were similarly reduced **(Figure 7D-E, M-N)**, suggesting dampened thrombus maturation and neutrophil activation. PI⁺ necrotic cell burden was also significantly lowered, supporting a protective effect against microvascular damage **(Figure 7O-P)**. Functionally, ABR-238901 substantially restored capillary perfusion, although pockets of reduced perfusion and vascular leakage remained detectable **(Figure 7P-Q)**. The treatment also led to a marked improvement in pulmonary perfusion, with flux values approaching, but not fully reaching, those of sham mice **(Figure 7R-S)**. Endothelial and epithelial oxidative stress markers were also reduced by treatment **(Figure 7T-U)**, as was the expression of VCAM-1 and E-selectin **(Figure 7V-W)**, indicating dampened endothelial activation.

Finally, systemic cytokine profiling revealed that 12 of the 15 upregulated cytokines, including IL-6, IL-1β, TNF, IFN-γ, MCP-1, and GM-CSF, were significantly reduced by ABR-238901 treatment **(Supplementary Figure 3),** supporting a broad anti-inflammatory effect. These findings identify S100A8/A9 as a mechanistic driver of neutrophil- and NET-mediated pulmonary microvascular injury following MI and demonstrate that its inhibition is sufficient to restore immune and vascular homeostasis in the lung.

## Discussion

Our study demonstrates that acute myocardial injury not only damages the heart but also triggers remote organ damage and instigates a potent systemic thrombo-inflammatory response with a particular tropism for the pulmonary microcirculation. This is mediated predominantly by the neutrophil alarmin S100A8/A9 (calprotectin), which activates the pulmonary endothelium and recruits a barrage of pro-inflammatory neutrophils and other leukocytes, and promotes microthrombi and aggregates. In addition to NETosis, these events impede capillary flow and reduce pulmonary perfusion, ultimately leading to cellular necrosis **(Graphical Abstract)**. Notably, these pulmonary changes closely mirrored those observed in the infarcted myocardium and exceeded thrombo-inflammatory responses seen in other organs such as the kidney and liver. Short-term pharmacological inhibition of S100A8/A9 markedly attenuated these effects, reducing platelet-neutrophil aggregates, NET formation, fibrin deposition, and microvascular obstruction, while substantially restoring pulmonary perfusion. Together with previous publications demonstrating improved cardiac outcomes with ABR-238901 treatment, our findings highlight S100A8/A9 as a central mediator linking cardiac injury to distal pulmonary dysfunction and suggest that targeting this pathway can alleviate both lung and cardiac injury following MI.

### The susceptibility of the pulmonary microcirculation to injury post-MI

MI has long been recognised as a trigger of systemic inflammation, but its impact on remote organs such as the lung has been underappreciated. Clinically, a severe MI can precipitate ALI or even ARDS in the absence of heart failure or infection, implicating inflammation as a key driver [4, 5]. In our study, despite a systemic “cytokine storm” (15 pro-inflammatory cytokines increased post-MI) that could affect multiple organs, the lungs emerged as the principal initial target of the heart’s inflammatory output. First, due to coronary vascular anatomy, the pulmonary circulation is the first tissue bed exposed to cardiac-derived inflammatory mediators. Venous blood from the coronary circulation empties into the right atrium and is pumped directly into the pulmonary circulation. As a result, the lungs receive the highest concentration of cardiac-derived DAMPs, cytokines, and activated leukocytes immediately after MI [32, 33]. Neutrophils or platelet-neutrophil aggregates formed in the cardiac vasculature may physically lodge within the lung’s dense capillary network, a well-recognised phenomenon in other conditions such as venous thromboembolism, where emboli typically impact the lungs [34, 35].

Second, the pulmonary capillaries offer a vast surface area for leukocyte-endothelial interaction [36]. We observed strong upregulation of endothelial adhesion molecules (E-selectin, VCAM-1) in MI lungs, effectively “priming” the lung vasculature to capture circulating neutrophils. In contrast, the liver, despite receiving a dual blood supply from both the hepatic artery and the portal vein, showed minimal neutrophil recruitment or S100A9 deposition. This suggests that factors beyond blood flow alone, such as endothelial activation state or local immune context, determine susceptibility to thrombo-inflammatory injury. The kidneys, which develop AKI in around 20% of MI hospitalisations (a known predictor of worse outcome), did exhibit injury signals, notably oxidative DNA damage, but neutrophil and platelet infiltration in the kidneys were much less pronounced than in the lungs [6]. It is possible that although the renal microvasculature is highly perfused, it is less predisposed to neutrophil trapping unless prolonged hypoperfusion or specific chemokine gradients are present [37]. The lung’s unique immunological environment, with constant exposure to circulating mediators and a primed endothelium, may further amplify thrombo-inflammatory signalling once triggered. Our comparative analysis highlights the organ selectivity of post-MI inflammatory injury and underscores that the cardiopulmonary axis is a particularly sensitive and vulnerable pathway following MI, akin to the well-established cardiorenal axis.

### Neutrophil- and NET-driven pulmonary disturbances after MI

With the pulmonary endothelium primed to capture circulating neutrophils and monocytes, we observed a rapid wave of immune and thrombotic activity targeting the lungs following myocardial reperfusion. The magnitude of these thrombo-inflammatory disturbances was comparable, if not equal, to that observed in the infarcted myocardium, underscoring the lung’s heightened vulnerability to MI-induced systemic inflammation.

This pattern aligns with a growing body of literature documenting neutrophil- and NET-mediated injury during post-infarction immune responses [38, 39]. Activated neutrophils release NETs, which drive tissue damage through extracellular DNA and DAMPs such as S100A8/A9. These mediators are released from dying cardiomyocytes and activated myeloid cells within hours of MI, serving as potent chemoattractants that amplify neutrophil recruitment and endothelial activation across distant vascular beds [40]; here we show that this also occurs within the distant pulmonary vasculature. NETosis, fuelled by this neutrophil activation, has been implicated not only in myocardial reperfusion injury itself but also in circulating microvascular events beyond the heart. In experimental models of ischaemia-reperfusion, NETs orchestrate remote organ dysfunction, including lung injury, via platelet activation and extracellular DNA-mediated thrombo-inflammation [41]. Human studies similarly show that circulating NET components correlate with MI severity, infarct size, and microvascular flow impairment [42].

In our study, we directly visualised NET formation in the post-MI lung microvasculature, observing a dramatic increase in NETs and fibrin deposition, confirming that neutrophils were not only recruited in large numbers but also actively participated in pro-thrombotic clot formation. These findings echo prior reports of NET-mediated thrombo-inflammation in ARDS, sepsis, and acute MI; NETs present in coronary thrombi can initiate intravascular coagulation [43, 44]. Our results suggest that a similar process unfolds remotely in the pulmonary vasculature following MI. The extensive co-localisation of NETs with platelets and fibrin implies a self-amplifying cycle in which NETosis triggers platelet activation and coagulation cascades, leading to microthrombus formation and further neutrophil trapping. This feed-forward loop of inflammation and thrombosis can severely compromise capillary blood flow, as shown by the steep decline in pulmonary perfusion we documented shortly after reperfusion. Regions of impaired perfusion corresponded to areas of necrosis, indicating that microvascular occlusions translated into lung injury. Such damage resembles that seen in diffuse alveolar injury associated with ARDS, although here it occurs in the absence of infection, reflecting a sterile inflammatory pathology [45, 46].

At the cellular level, we detected significant increases in 8-OHdG, a marker of oxidative DNA damage, in both endothelial and epithelial compartments. This suggests that pulmonary injury extends beyond the vasculature into the parenchyma, potentially compromising alveolar integrity and gas-exchange function. Together, our results align with clinical reports suggesting that patients with recent MI are at increased risk of non-cardiogenic pulmonary complications, including impaired gas exchange, pulmonary inflammation, and radiological signs of lung injury, phenomena often dismissed as subclinical or attributed solely to haemodynamic overload [47, 48].

This sterile, MI-induced lung injury appears to parallel neutrophil-driven lung inflammation observed in other contexts, suggesting a shared pathogenic axis. In viral exacerbations of asthma [49, 50] and COPD [22], for example, excessive NET formation has been shown to promote neutrophilic inflammation, microvascular dysfunction, and tissue damage. In both conditions and our MI model, neutrophil activation and NETosis initiate a cascade of thrombo-inflammation that disrupts capillary perfusion and injures the lung. Thus, myocardial infarction, even in the absence of infection, can provoke lung injury via systemic immune crosstalk, representing immune-mediated remote organ damage rather than a mere haemodynamic consequence.

### S100A8/A9 as a neutrophil-derived and NET-associated inflammatory driver

Among the most potent neutrophil-derived and NET-associated inflammatory mediators is the S100A8/A9 complex (calprotectin), which is abundantly expressed in activated neutrophils and monocytes. Functioning as a DAMP, S100A8/A9 amplifies leukocyte-driven inflammation and also modulates platelet activation and coagulation, positioning it at the centre of thrombo-inflammation [51]. When secreted extracellularly, S100A8/A9 acts as a cytokine and a chemokine, promoting endothelial and neutrophil activation. Notably, during NETosis approximately 40% of S100A8/A9 becomes bound to the NETs, while the remainder circulates as soluble protein or in microvesicles [52]. Recent mechanistic studies have established that S100A8/A9 is a potent driver of inflammation during MI [17, 51], and our work now connects this alarmin to post-MI pulmonary injury.

In line with this, we observed that S100A8/A9 is rapidly released during MI, with marked upregulation in the lungs and plasma of MI patients and in mice, reaching levels comparable to those seen in COPD patients. Although direct comparison with classical cytokines such as IL-6, IL-1β, TNF, and IFN-γ is limited by differences in baseline abundance and potency, the pronounced induction of S100A8/A9 highlights its potential as a dominant mediator of the pulmonary response, driving neutrophil recruitment and endothelial activation.

S100A8/A9’s pathological significance in MI is supported by clinical studies linking high plasma levels during MI with worse left ventricular function and a higher risk of heart failure [14, 17]. Our findings extend this correlation to the pulmonary complications of MI, showing strong co-localisation of S100A9 with infiltrating neutrophils in the pulmonary microvasculature, as well as abundant extracellular release in areas of intense neutrophil accumulation. This resembles observations in sepsis and ARDS, where neutrophil-derived DAMPs and NETs drive local tissue injury [51].

Mechanistically, extracellular S100A8/A9 acts as a cytokine to activate endothelial cells and platelets and amplify neutrophil adhesion and NETosis, primarily through Toll-like receptor 4 (TLR4) and the receptor for advanced glycation end-products (RAGE) on both neutrophils and platelets [53]. In models of pneumonia, S100A8/A9 released during inflammation stimulates neutrophil recruitment and facilitates platelet-neutrophil interactions in the lung, largely through TLR4/RAGE signalling, driving microvascular trapping [54]. In our MI model, we saw frequent platelet-neutrophil aggregates within lung capillaries, consistent with an S100A8/A9-driven recruitment axis that amplifies thrombo-inflammation and promotes the formation of these pathogenic aggregates. Importantly, recent work has shown that neutrophils rapidly release S100A8/A9 upon E-selectin engagement, via NLRP3 inflammasome activation and gasdermin D-mediated membrane pores, a mechanism that can occur independently of full NETosis [55, 56]. This process may further explain the swift extracellular alarmin release in early reperfusion lung injury.

Interestingly, our human lung samples from patients up to eight months post-MI still showed elevated S100A8/A9 expression and microvascular perturbations, despite the absence of overt acute inflammation. This persistence mirrors other inflammatory conditions, such as post-ICU sepsis or COVID-19, where circulating S100A8/A9 remains elevated for months and correlates with ongoing lung dysfunction [57]. While our study focused on acute mechanisms, these observations suggest S100A8/A9-driven pathways could contribute to longer-term pulmonary changes after MI. The precise reasons why S100A8/A9 remains elevated chronically are not yet clear, but possibilities include sustained endothelial priming, prolonged low-level release from activated myeloid cells, or other unresolved inflammatory signals. Notably, the extent of S100A8/A9 induction and microvascular disturbance we observed post-MI closely resembled those seen in COPD, one of the world’s leading causes of death, further highlighting the pathological significance of this response. Together, these data support S100A8/A9 as a central orchestrator of remote lung injury after MI, linking cardiac damage to thrombo-inflammation in the pulmonary circulation. Its rapid release and broad activity, promoting neutrophil recruitment, NETosis, platelet adhesion, and endothelial activation, make it a compelling target for therapeutic intervention to preserve both cardiac and pulmonary microvascular integrity following MI.

### Therapeutic impact of inhibiting neutrophil-derived and NET-associated S100A8/A9

A principal outcome of our study is the evidence that inhibiting S100A8/A9 can effectively disrupt the cascade of remote lung injury after MI. Short-term treatment with the small-molecule inhibitor ABR-238901 resulted in marked protection of the pulmonary microcirculation. We observed significantly fewer neutrophils and platelets adhering in lung capillaries, minimal NET formation, reduced fibrin clots, and preserved capillary perfusion in ABR-treated mice. This immune rebalancing likely reflects, at least in part, the drug’s acute effects on neutrophil activation, with additional contributions from its known effects on haematopoiesis and reduced neutrophil recruitment [51, 58].

Previous work in mouse MI models showed that S100A9 inhibition during the acute phase attenuates the systemic inflammatory response by inhibiting haematopoietic stem cell proliferation and myeloid cell egress from the bone marrow [17]. Consequently, fewer neutrophils and inflammatory monocytes flood the circulation and infiltrate the heart, fostering a more anti-inflammatory environment and limiting cardiac damage. Similar protective effects were observed in global S100A9-deficient mice, which also lack S100A8, in which infarct size, cardiomyocyte death, and adverse remodelling were reduced [51]. Our findings extend these observations to the lungs: by tempering the leukocyte surge, S100A8/A9 inhibition prevents the downstream platelet-neutrophil interactions and endothelial injury that cause microvascular obstruction. Put simply, S100A8/A9 inhibition dampens the upstream triggers of thrombo-inflammation, yielding dual benefits, reducing myocardial injury, and, as we demonstrate here, protecting the lung’s fragile microvasculature [17, 58]. This dual action is particularly noteworthy given that multi-organ dysfunction portends worse outcomes after MI.

Therapeutically, our data reinforce the case for S100A8/A9 as a promising target in acute coronary syndromes: not only might its inhibition improve cardiac function and reduce heart failure incidence, but it could also avert complications such as acute lung injury. While other anti-inflammatory strategies (e.g. IL-1 inhibitors or colchicine) have been tested in MI to prevent adverse cardiac remodelling, they generally do not address the immediate microvascular damage in remote organs [59, 60]. Targeting a neutrophil/monocyte-derived alarmin such as S100A8/A9 represents a novel approach to blunt the initial “cytokine storm” and thrombo-inflammatory cascade at its origin.

## Conclusion

Our findings highlight a novel facet of heart-lung crosstalk in which an MI unleashes a neutrophil- and S100A8/A9-mediated thrombo-inflammatory cascade in the lungs, compromising the pulmonary microcirculation. We demonstrate that this process closely mirrors the inflammatory and microvascular injury within the infarcted myocardium and, notably, persists beyond the acute phase. This pathogenic pathway, a sterile inflammatory process reminiscent of sepsis-induced lung injury, may underlie the respiratory complications observed after infarction. Importantly, we show that therapeutic targeting of a single upstream mediator, S100A8/A9, is sufficient to markedly reduce pulmonary thrombo-inflammation and preserve lung perfusion in our preclinical models. These results underscore the translational potential of S100A8/A9 inhibition as a precise, organ-protective strategy that not only preserves myocardial integrity but also mitigates secondary pulmonary and systemic injury. Our findings support the rationale for clinical studies to evaluate S100A8/A9 inhibition, with a focus on optimising timing, assessing safety, and evaluating long-term efficacy in patients.

## NONSTANDARD ABBREVIATIONS AND ACRONYMS

Abbreviation: Full Term
MI: Myocardial infarction
IRI: Ischaemia-reperfusion injury
ALI: Acute lung injury
ARDS: Acute respiratory distress syndrome
DAMP: Damage-associated molecular pattern
NET: Neutrophil extracellular trap
COPD: Chronic obstructive pulmonary disease
FCD: Functional capillary density
PI: Propidium iodide
8-OHdG: 8-hydroxy-2’-deoxyguanosine
LSCI: Laser speckle contrast imaging
VCAM-1: Vascular cell adhesion molecule-1

## Authorship Contribution

D.L., O.K., and E.M. acquired and analysed the data; B.G. and M.J. acquired the data; C.C., E.J., K.S., E.A., and N.K. collected patient samples; B.N. performed surgical lung resections; R.M., D.R.T., A.Sc., and E.S. assisted with obtaining clinical samples and contributed funding, resources, and data interpretation; D.K. and H.M. contributed to data interpretation and provided funding and resources; J.R. contributed to conceptualisation and data interpretation, and provided funding and resources; Al.S. provided the inhibitor and contributed to data interpretation; S.J. and A.Si. provided samples, contributed to data interpretation, and provided funding and resources; J.E.A. obtained funding, designed the experiments, acquired, analysed, and interpreted the data, and drafted the manuscript. All authors reviewed and approved the final version of the manuscript.

## Acknowledgments

We acknowledge the support of the Intravital Imaging Suite, RRID:SCR_027189, Technology Hub Facilities at the College of Medicine and Health, University of Birmingham, for providing access to equipment and technical expertise. We also thank the Biomedical Services Unit (BMSU) at the University of Birmingham for their continued support and expertise during the *in vivo* aspects of this study.

